# Oxytocin regulation of social transmission of fear in zebrafish reveals its evolutionary conserved role in emotional contagion

**DOI:** 10.1101/2021.10.06.463413

**Authors:** Ibukun Akinrinade, Kyriacos Kareklas, Michael Gliksberg, Giovanni Petri, Gil Levkowitz, Rui F. Oliveira

## Abstract

Emotional contagion is the most ancestral form of empathy that relies on simple perception-action mechanisms, on top of which more complex forms of empathic behaviors, such as consolation and helping, have evolved. Here we tested to what extent the proximate mechanisms of emotional contagion are evolutionary conserved by assessing the role of oxytocin, known to regulate empathic behaviors in mammals, in social fear contagion in zebrafish, which represents an evolutionary divergent line to that of tetrapods, within vertebrates. Using mutants for the ligand of the fish oxytocin nonapeptide and both of its receptors in zebrafish we showed that oxytocin is necessary for observer zebrafish to copy the distressed behavior of conspecific demonstrators. Exogeneous administration of oxytocin to the ligand mutant rescued the ability of observers to express social fear transmission, indicating that oxytocin is not only necessary but also sufficient for emotional contagion. The brain regions in the ventral telencephalon that are associated with emotional contagion in zebrafish are homologous to those known to be involved in the same process in rodents (e.g. striatum, lateral septum), and receive direct projections from oxytocinergic neurons located in the pre-optic area. Finally, we ruled out the hypothesis that social transmission of fear in zebrafish merely relies on behavior contagion by motor imitation, and we showed that it rather relies on emotion discrimination. Together our results support an evolutionary conserved role for oxytocin as a key regulator of basic empathic behaviors across vertebrates.

**One-Sentence Summary:** Oxytocin is necessary and sufficient for social fear contagion in zebrafish supporting an evolutionary conserved role for oxytocin in emotional contagion among vertebrates.

## Main Text

Emotional contagion, described as the ability to match the emotional state of another individual, has been considered the most ancestral form of empathy, on top of which more complex forms of empathic behaviors, such as consolation and helping, have evolved in species endowed with more complex cognitive abilities (e.g. rodents, elephants, dolphins, primates [1-3]). Emotional contagion relies on simple perception-action mechanisms and provides important adaptive advantages to social living species [1]. It enhances social cohesion and the establishment of social bonds and promotes the rapid spread of fear among group members once a threat is detected (e.g. predators), hence allowing individuals to survive potential dangers without directly experiencing them [3]. Therefore, emotional contagion is expected to be phylogenetically ancient, being present even in species with less elaborate social cognition. Indeed, social contagion of fear has been recently described in zebrafish (*Danio rerio*), consisting the transmission of distress behavior to observers and increases in observer cortisol levels similar to those of target individuals [4-6]. Moreover, behavioral responses were influenced by familiarity, with familiar distressed target fish eliciting stronger distress responses in observers [5]. The face validity of this transmission of distress behavior as emotional contagion can also be argued by comparison with mammalian models of emotional contagion, such as facial expressions in orangutans [7] and freezing in rodents [8]. However, to what extent the social contagion of fear observed in fish and in mammals is homologous, or represents a case of convergent evolution, remains an open question. To disentangle these two hypotheses, we investigated if emotional contagion in zebrafish shares the same proximate mechanism that have been described for mammals. We focused on the oxytocin signaling system since nonapeptides of the oxytocin family are evolutionary conserved across vertebrates [9] and have been implicated in the regulation of emotional contagion in rodents [10, 11] and fear recognition in humans [12, 13].

Here we have used zebrafish mutant lines for the ligand (*oxt*) and the two receptors (*oxtr* and *oxtrl*) of the zebrafish oxytocin nonapeptide (CYISNCPIG-NH_2_, aka isotocin [9]) to assess the role of oxytocin on fear contagion. In zebrafish injured individuals release an alarm substance from their skin into the water, which is detected through olfaction eliciting a distress response, which consists of erratic movement followed by freezing behavior [14]. The sight of conspecifics in distress also elicits the expression of this response in observers, indicating the occurrence of fear contagion in zebrafish [4-6] (Fig. 1). Therefore, we have used an experimental paradigm in which a naïve observer fish watches a conspecific shoal in a neighboring tank (i.e. without chemical communication), to which we have administered either water (control) or the alarm substance (Fig. 1A). Given that freezing behavior was a more consistent distress response in wild type than erratic movement (Fig. 1B-K), we have used it as a read-out for fear contagion. Observer individuals of all wild type control lines (i.e. *oxt*^+/+^, *oxtr*^+/+^ and *oxtrl*^+/+^) significantly increased their freezing behavior when exposed to a distressed shoal. In contrast, observer individuals of all the mutant lines (i.e. *oxt*^-/-^, *oxtr*^-/-^ and *oxtrl*^-/-^) failed to significantly increase their freezing behavior when exposed to a distressed shoal (Fig. 1C, E, G). This indicates that oxytocin signaling is necessary for fear contagion in zebrafish. Moreover, we have administered exogenous oxytocin to the ligand mutants and their controls to assess if it could recue the fear contagion phenotype. We have also injected another group of ligand mutants and their respective control with the vehicle solution to control for putative stressful effects of the injection. Mutants injected with oxytocin also increased significantly their freezing behavior when exposed to distressed conspecifics, indicating that oxytocin is both necessary and sufficient for emotional contagion (Fig. 1E, G).

**Fig 1.**
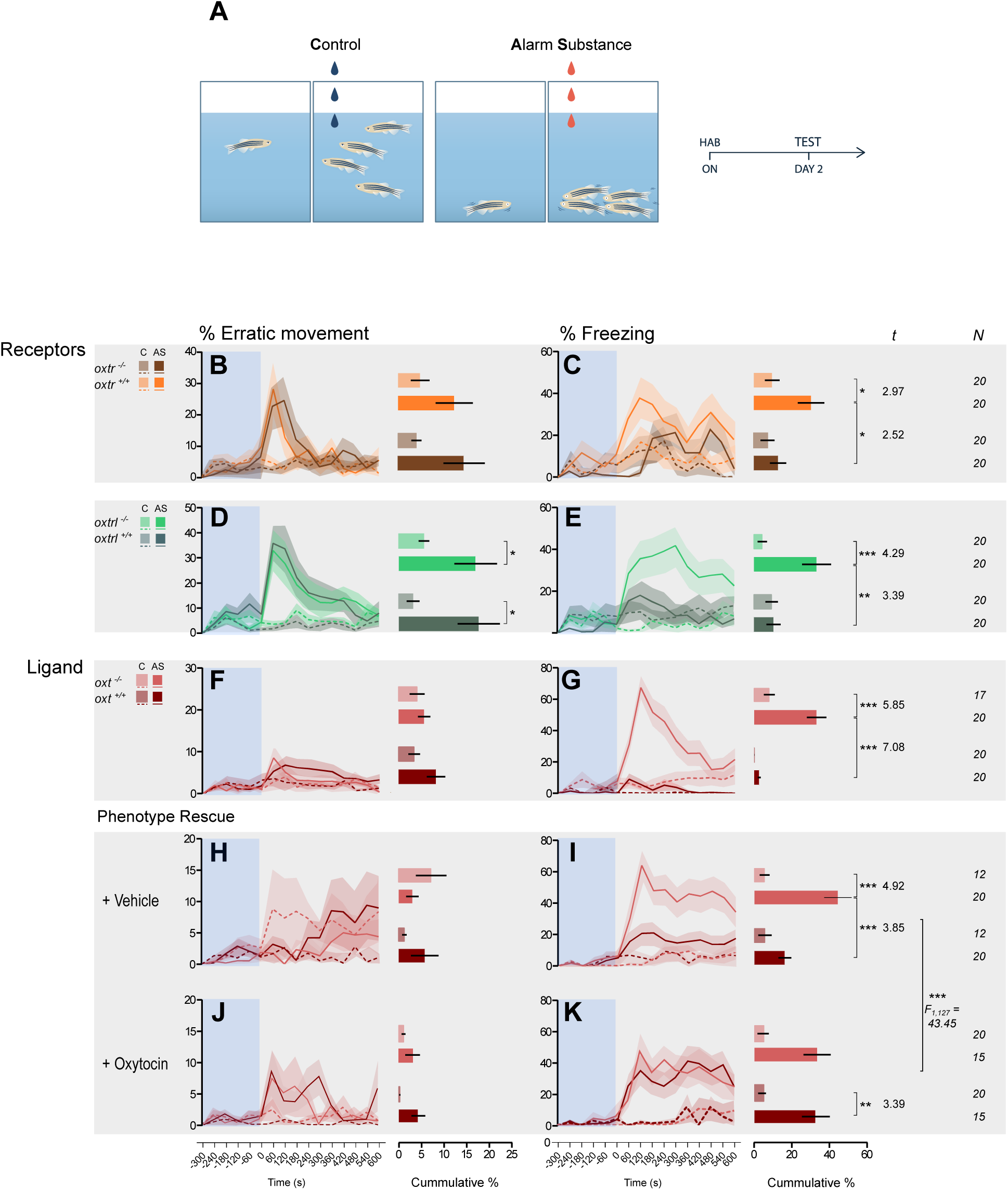
Oxytocin effects on the social transmission of distress. (**A**) Schematic and schedule (HAB, habituation; ON, overnight) of social contagion of fear paradigm. Droplets represent administration of vehicle (blue) and alarm substance (AS; red) to control and experimental groups. (**B-K**) Left panels: temporal dynamics of freezing and erratic movement response across treatments for mutant oxtr^(-/-)^,oxtrl^(-/-)^,oxt^(-/-)^ and ligand rescue with intraperitoneal (i.p) oxytocin (OXT) to oxt^(-/-)^ fish. Shaded area indicates time before AS or vehicle administration. Right panel: percentage of freezing and erratic movement (mean±SEM) after vehicle and AS administration. [*p < 0.05, **p < 0.01, ***p < 0.001]

In order to characterize the neural circuits associated with emotional contagion in zebrafish we examined the expression of a neuronal activity marker, phospho-S6 ribosomal protein (pS6), across a set of forebrain and midbrain areas known to be involved in social decision-making across vertebrates (aka social decision-making network, [15]). Two of these brain regions (Fig. 2A), the ventral nucleus of the ventral telencephalic area (Vv), which is a putative homologue of the mammalian lateral septum, and the central nucleus of the ventral telencephalic area (Vc), which is a is a putative homologue of the mammalian striatum, show a significant decrease in activity associated with the expression of freezing behavior in observer wild-types, and a significant increase in activity associated with the lack of distress response in the *oxtr* mutants (Fig. 2B, C). This result suggests that zebrafish distress behaviour during emotional contagion is mediated by decreases in inhibitory cell activity in Vv and Vc and that mutants have an overactivation of these cells that prevents the expression of this behavior. Oxytocin regulation of these ventral forebrain areas is supported by the occurrence of projections from pre-optic area oxytocin neurons to these areas (Fig.2C), as well as by the expression of both zebrafish oxytocin receptors (*oxtr, oxtrl*; Fig S1) in these areas. Interestingly, both receptors are also expressed across most nodes of the social decision-making network, but the expression of the secondary receptor (*oxtrl*) is distinctly lower and less widespread (Fig. S1). Remarkably, the expression, recognition and sharing of emotions in humans also relies on the activity in these forebrain areas and their regulation by oxytocin, even if higher order empathic functions are dependent of neocortical circuits [16-19]. In particular, the striatum is implicated in downstream other-oriented processes of emotion recognition, such as sympathy and compassion [16], whereas the lateral septum is an area of functional connection between the reward system and the social behavior network, and is involved in the regulation of emotional expressions [17].

**Fig 2.**
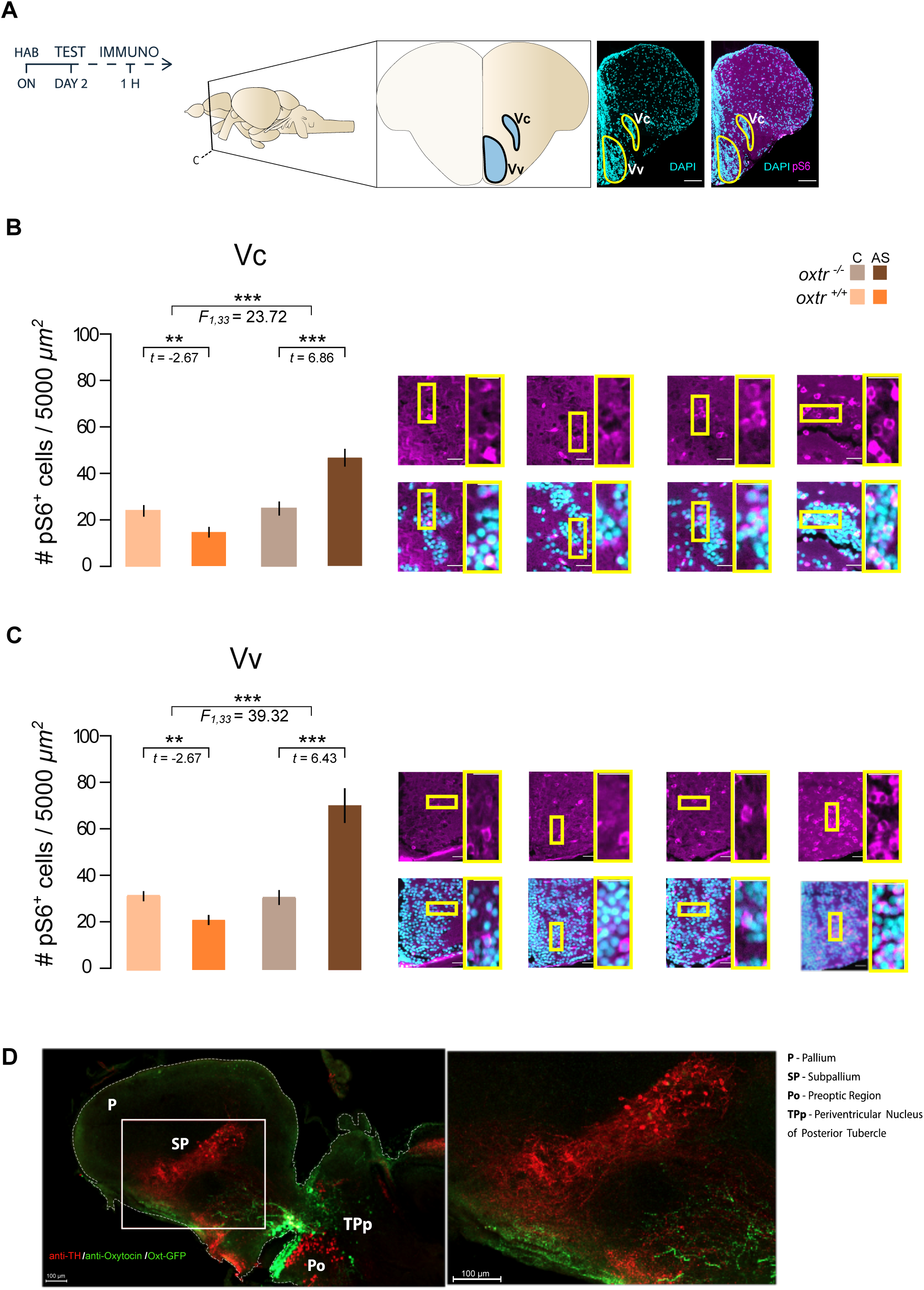
Oxytocin receptor deletion alters nodal neuronal activation. (**A**) Schedule of behavioural assay and immuno-staining and anatomical localization of the two brain areas that respond to the social contagion of fear: ventral nucleus of the ventral telencephalon (Vv) and the central nucleus of the ventral telencephalon (Vc), with representative hemi-coronal sections identified by DAPI (cyan) and patterns of neuronal shown by pS6 (magenta) via immunostaining. (**B, C**) Quantification of the density (cells/5000µm2) of pS6 positive cells in each brain area (identified above the graph); panels show representative examples (left to right: wild-type control, wild-type alarm, mutant control, mutant alarm; scale= 20 µm). The result is shown as Mean±SEM. [*p < 0.05, **p < 0.01, *** p < 0.001]. (**D**) Representative example of sagittal brain slice (confocal maximum intensity z-stack) showing immunostained *oxt* positive neuronal fibers (green) projecting to the Vv in the subpallium of adult zebrafish highlighted with the red square with TH cell groups (red) also projecting to the Vv.

In order to study the patterns of functional connectivity across the social decision-making network, we also constructed networks representing the co-activation patterns for each treatment, with positive and negative correlations between nodes respectively indicating excitatory and inhibitory patterns (Fig. 3). Under control conditions wild-types and *oxtr* mutants show differences in the network distribution of excitation (*KS* = 0.23, *p* < 10^−4^) and inhibition (*KS* = 0.55, *p* < 10^−6^), but negligible differences in average signals (only inhibition: *U* = 3492, *p* < 10^−5^, Cohen’s *d* =0.18). However, when exposed to distressed others, *oxtr* mutants exhibited both higher average excitation (*U* = 5744, *p* < 10^−6^, Cohen’s *d* = 0.67) and inhibition (*U* = 1972, *p* < 10^−6^, Cohen’s *d* =0.68), as well as differences in distribution (excitation: *KS* = 0.49, p<10^−6^; inhibition: *KS* = 0.73, p<10^−6^). Moreover, while wild types show no changes when exposed to distressed conspecifics, *oxtr* mutants show large increases in excitation (*U* = 5680, *p* < 10^−6^, Cohen’s *d* = 0.49) and marginal differences in inhibition (*U* = 1366, *p* = 0.01, Cohen’s *d* = 0.06). The two genotypes also show differences in both the distribution of network excitation (*KS* = 0.38, *p* < 10^−6^) and inhibition (*KS* = 0.50, *p* < 10^−4^), with mutants showing a segregation of patterns into smaller modules and wild types an integration of patterns across the network (Fig. 3). In summary, the absence of emotional contagion in *oxtr* mutants is paralleled by a segregated pattern of functional connectivity with significantly larger excitation in their brain network than wild types that display the socially transmitted distress behavior. In order to further capture changes in the structure of the brain networks between treatments, we have also computed the centrality (aka strength, weighted degree of the full correlation matrix, Fig. S2) of nodes and rank them in descending order (Table S1). We find that the node centrality rankings change between conditions quite radically, since there was only one positive significant correlation in wild type individuals in the control vs. exposed to distressed conspecifics (Kendall’s *τ* = 0.36, *p* = 0.03). This implies that the hub structure and relevance changes radically between brain networks of wild type and *oxtr* mutants, as well as between the control and exposure to distressed conspecifics treatments in the *oxtr* mutant group itself. Finally, we have used a community detection technique to look for conserved network submodules between conditions and groups. However, we found that the modular structure of the network reconfigures radically across the various treatments, and the only conserved submodule identified (at *z* > 3, *p* < 0.01) is shared between wild types and *oxtr* mutants when exposed to distressed conspecifics, and is composed by the dorsal and ventral habenula, the posterior POA (PPp) and PM. Considering that the habenular nuclei have been implicated in fear responses in zebrafish, and that the POA is known to respond to the alarm substance, we hypothesize that this conserved module is involved in processing the fear stimulation in both groups. Interestingly, this conserved submodule is isolated in the *oxtr* mutants, while it is integrated in a larger module (i.e. D, VS, DL, PM, PPP, HAD, HAV, HV) in wild types, pointing to a larger brain functional integration in wild types than in *oxtr* mutants when exposed to distressed conspecifics.

**Fig 3.**
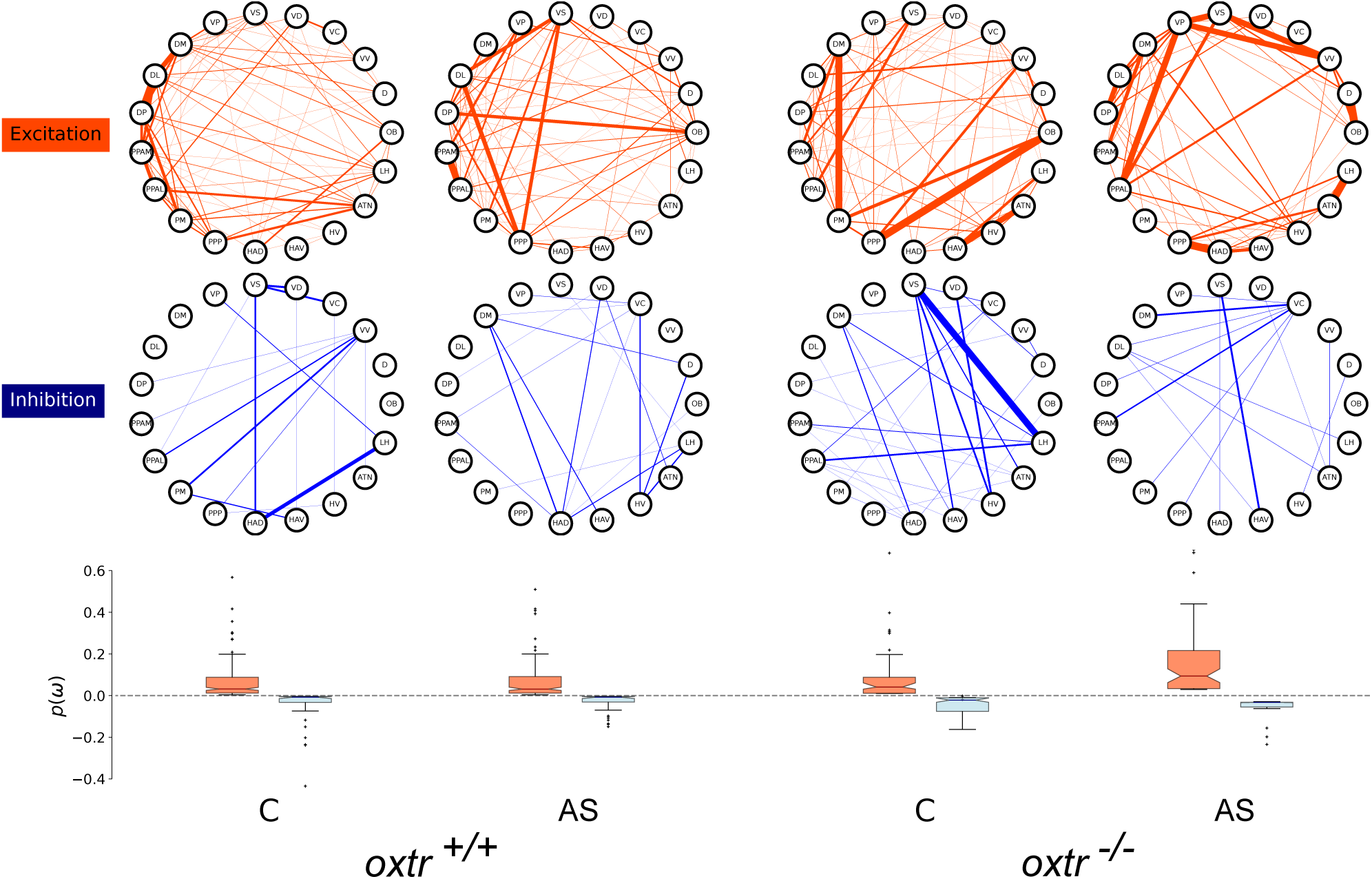
Network analysis of co-activation patterns in the social decision-making network. Nodes represent different regions and edges the relationship between them, where greater correlation values are represented by greater thickness. Networks were tested for both excitatory and inhibitory distributions across genotypes and treatments, and for computed average levels of each [probability in sample space, *p(ω)*].

The observed social transmission of fear in zebrafish can be regarded simply as behavior contagion based on motor imitation, or as emotional contagion, which requires the recognition of the demonstrator’s state (i.e. emotion) which triggers an automatic representation of the same state in the observer, causing an equivalent expression of behavior [20] Although the available evidence supports the latter, namely the fact that zebrafish observers not only express similar behaviors as demonstrators but also match their cortisol levels [4], which can be taken as a measure of their internal state, we have decided to further rule out the simpler explanation of behavioral contagion using a video-playback experimental paradigm. Observer fish were exposed to two LCD monitors in which previously recorded videos were played-back. In one of the videos the demonstrator fish swims in a neutral state, whereas in the other video the same demonstrator fish periodically exhibits distress behavior, which consists of three bouts of an erratic and freezing repertoire. After this observation phase, observers were exposed to two similar videos with the conspecific swimming in the neutral state (Fig. 4A). Therefore, if observers discriminate between the two videos in the test phase, this cannot be explained by behavioral contagion because in both videos the conspecific is not expressing distress behavior during this stage of the experiment, and they have to rely on the distress behavior observed in one of the videos in the previous observation stage of the experiment. During the observation phase we found that simultaneous video playbacks of distressed and neutral behavior in the same demonstrator, which targeted state valence alone, elicited orienting preferences in observers for both erratic movement and freezing (Fig. S3). Thus, overall attention shifted to the distressed behavior and not the level of movement. Notably, these body orientation preferences were not different between any of the oxytocin mutants and their respective wild type controls (Fig. 4B-E). In contrast, and in agreement with the experiment with live demonstrators, during the observation phase oxytocin mutants (*oxt, oxtr, oxtrl*) fail to mimic the distress behavior of the demonstrators, whereas their respective wild-type controls do respond with distress behavior, and the administration of oxytocin to the ligand mutant (*oxt*) rescues the expression of the fear contagion (Fig. 4F-I), but peaks and troughs in speed and angular velocity rarely reached the levels of demonstrators (Fig. S4). Therefore, the video-playback experiment replicated the results of the live demonstrator experiment regarding the necessary and sufficient role of oxytocin for fear contagion and further suggested that attention is not moderating these effects of oxytocin. The results of the test phase of the video-playback experiment show that observer wild-type zebrafish can discriminate demonstrators based on their previous behavior (distressed vs. neutral) during the observation phase, but oxytocin mutants (*oxt, oxtr, oxtrl*) fail to do so, and that the discrimination can be rescued in ligand mutants treated with exogenous oxytocin (Fig. 4J-Q). Interestingly, this discrimination is expressed by a motivation in wild type observers to more readily approach and prefer being near the demonstrator that they had previously observed in a distress state, whereas oxytocin mutants do not express a motivation to approach the previously distressed demonstrator (Fig. 4 J-M), and prefer being near the demonstrator that was in a neutral state in the observation phase (Fig. 4 N-Q). Taken together, the results of the video-playback experiment indicate that oxytocin is necessary and sufficient for zebrafish to discriminate between conspecifics only based on their altered states, suggesting the occurrence of the oxytocin-dependent ability to discriminate distressed from neutral behavior in conspecifics (aka emotion recognition [21, 22]) in zebrafish. This is in line with responses in humans and mammals that implicate the recognition of a fearful state in others and not simply their behavior [23, 24]. Moreover, because distressed behavior in zebrafish signals local predation risk [14], our findings also show that oxytocin promotes interaction with distressed others despite heightened local risk. Such other-oriented acts that involve individual costs and benefits for others are typically referred to as prosociality, which is well-defined in mammals where it is also regulated by oxytocin [2, 20 - 22]. In this case the benefit for the receiver can be due to the buffering of distress in the presence of others, which has been also described in zebrafish [25]. Thus, approaching and interacting with a distressed individual may prove to be a prosocial behavior in zebrafish, but further evidence is needed.

**Fig. 4.**
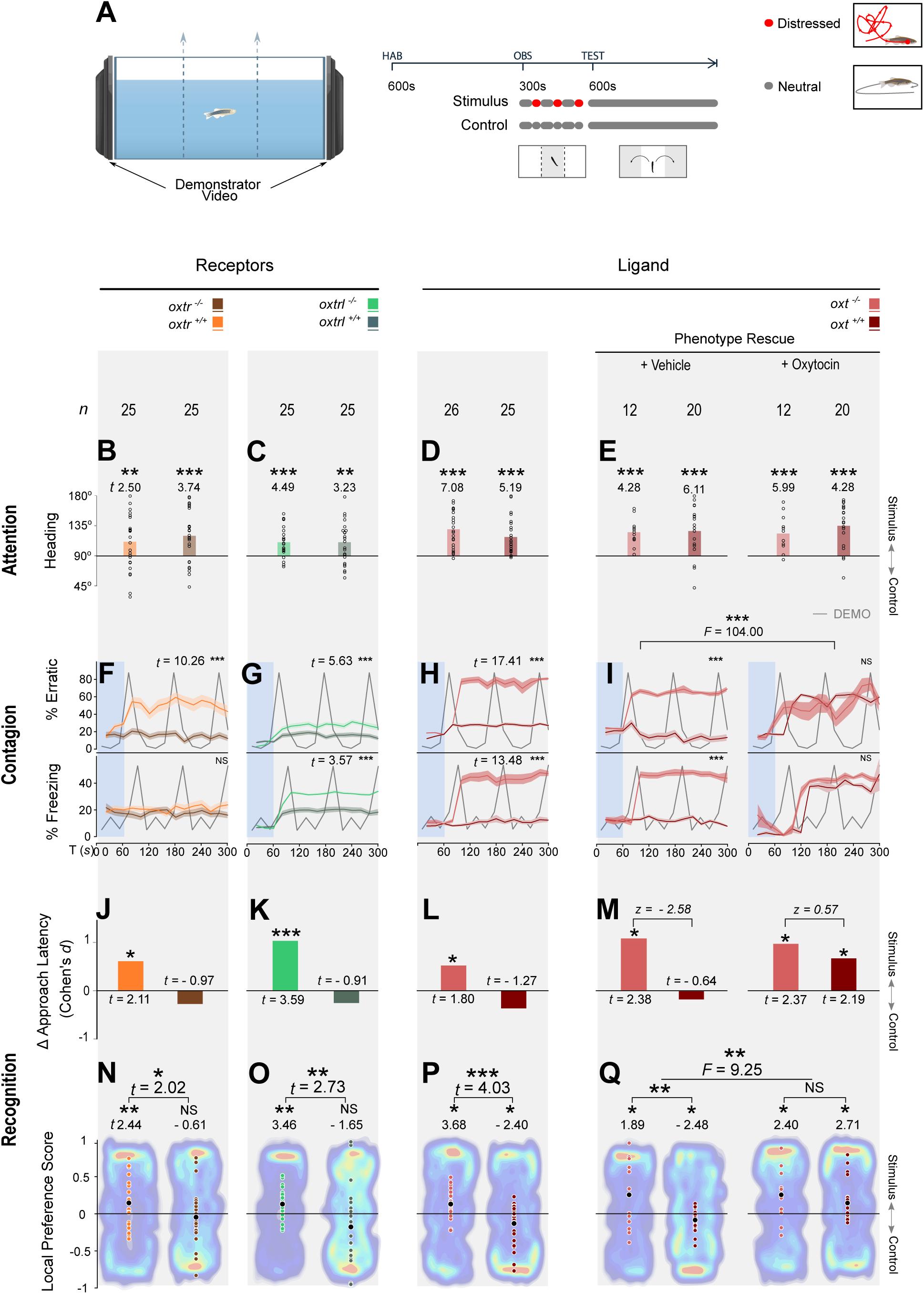
Content validity of fear transmission by state recognition. (**A**) Video playback tests enabled the controlled assessment of stimulus versus state recognition across two experimental phases: a 5 min observation of two conflicting videos presenting the same demonstrator in either neutral (control) or periodically distressed (stimulus) state; a 10 min local-preference test while both videos displayed the demonstrator in a neutral state. During observation, (**B-E**) attention was measured by the absolute heading towards the stimulus video (0 – 180 °; 1-sample *t*-tests, *µ* ≠ 90°) and (**F-H**) temporal changes in the proportion time erratic and freezing following analogous behaviour in the stimulus video were compared between genotype and treatment (LMMs, full factorial). During tests, (**J-M**) differences in latency to first approach between the stimulus and control location were tested (Welch’s 2-sample *t*-tests; effect-size comparisons: *z* tests, *d*_*1*_ ≠ *d*_*2*_, |*z*| ≥ 1.96 at *α* = 0.05 two-sided) and (**N-Q**) local preference scores calculated based on cumulative durations (1-sample *t*-tests, *µ* ≠ 0; genotypic comparisons: Welch’s 2-sample *t*-tests; genotype × treatment: two-way ANOVA with *post hoc* Fisher’s LSD). Heat maps are representative examples with the least deviation from the mean. [^NS^*P* > 0.05, \**P* < 0.05, \*\**P* < 0.01, \*\*\**P* < 0.001]

The oxytocin regulation of fear contagion in zebrafish described here supports its evolutionary conserved role in emotional contagion, given its similar effects in mammals, where exogeneous administration of oxytocin increases observational fear responses [10, 11]. Furthermore, both in zebrafish and in rodents oxytocin also regulates emotion recognition [23, 26], which is the cognitive basis for emotion contagion. Therefore, it is plausible that oxytocin has been recruited early in the evolution of nonapeptides to regulate ancestral empathic behaviors in group living species, and that it has been evolutionary co-opted to regulate more complex empathic behaviors, such as consolation and helping [24, 27-30], in species with more complex cognitive abilities. From a translational research perspective our results provide content and construct validity to a phylogenetically distant model of emotional contagion.

