## Supplemental methods, table and figures for "Oxytocin regulation of social transmission of fear in zebrafish reveals its evolutionary conserved role in emotional contagion"

**This PDF file includes:**

Methods

Table S1

Figs. S1 to S5

### Methods

Animals, housing and husbandry

We used naïve adult wild-type (WT) zebrafish, *Danio rerio,* from the *TU* strain (6-12 months), and adults of the transgenic *oxt*:EGFP reporter line from a *TL*-mixed background for characterizing the expression of oxytocin receptors and oxytocin fibre projections respectively,. In addition, we used WT and genetically modified (GM) adults from lines of a mixed TL background for testing the effects of oxytocin function on the social transmission of distress and the recognition of fear. All fish were housed in groups at a density of 10/L in a recirculation life support system (Tecniplast) maintained at 28 °C, pH 7.0, conductivity 1000 $\mu$S/cm and 14 L:10D photoperiod. Feeding included a combination of live (*Paramecium caudatum* and *Artemia salina)* and dry food (Gemma). Husbandry and health maintenance protocols were followed as previously described [31], and fish were kept free from known pathogens via sentinel testing. All experiments were conducted in accordance with standard operating procedures of the Institutional Ethics Committee, assessed and monitored by the Animal Welfare Body, and licensed by the National Competent Authority (DGAV-Direcção Geral de Alimentação e Veterinária, Portugal) with the permit number 0421/000/000/2020.

Genetic line characterization: modification of oxytocin signaling

The *oxt* mutant line (ZFIN ID: ZDB-ALT-180904-7) is a functionally null line with a small deletion of 7 base pairs in Exon 2 following treatment with CRISPR1-oxt at the embryonic stage, described by Blechman et al. [32]. It is a frameshift mutation leading to disruption of the translational reading and abolishing the expression of the oxytocin neuropeptide, and thus overall oxytocin signaling. The *oxtr* mutant line (ZFIN ID: ZDB-ALT-190830–1) is a functionally null mutant line that has a small deletion of 1 base pair following treatment with TALEN1-oxtr at the embryonic stage. It is a frameshift mutation that abolishes the expression of oxytocin receptor 1. The characterization of the line has been further described by Nunes et al. [33]. The *oxtrl* transgenic line (ZFIN ID: ZDB-ALT-190819-1) has an insertion of a multi-frame stop cassette (83bp) at the ATG+260 position leading to a stop codon formation after the 89$^{th}$ amino acid, following CRISPR treatment at the embryonic stage, which abolishes the expression of the oxytocin receptor 2. Finally, the *oxt:EGFP* transgenic reporter line (ZFIN ID: ZDB-ALT-111103-1) was generated using the Tol2kit transposon-based vector system for encoding the *oxt* gene and report the endogenous expression of *oxt* mRNA and protein [32]. All genetic lines were generated at the Weizmann Institute of Science, Israel, by G. Levkowitz, and in collaboration with R. Nunes for the *oxtr* transgenic line and M. Gliksberg for the *oxtrl* line.

Genotyping

Genotyping was performed by PCR of the genomic region of interest from clipped fins, followed by sequencing [33]. We designed specific primer pairs to target the deletion sites of the ligand [*oxt* (NM_178291.2): 5` − AGACACAAACACTAAGTAA − 3` (forward), 5` − AGCAGACGGACAGCAGACACAGCA − 3` (reverse)] and receptors [*oxtr* (NM_001199370.1): 5‘ − TGCGCGAGGAAAACTAGTT − 3‘ (forward), 5‘ − AGCAGACACTCAGAATGGTCA − 3‘ (reverse); *oxtrl* (NM_001199369.1): 5‘ − TTTTACGCACAATGGAGAGCC − 3‘ (forward), 5` − AGCATGTAAGTGGACGCGAA − 3’ (reverse)].

Alarm substance extraction

Alarm substance was extracted by following approved procedures under institutional and project licenses. Briefly, 12 adult fish of either sex (to control for variations) were euthanised via rapid chilling, placed on a petri dish kept on ice and 15 superficial surgical-blade cuts were performed on either side of their trunk to induce the release of alarm substance from the club cells. Cuts were then washed with 50ml of distilled water and filtered with a 240mm filter paper (VWR cat no. 516-0287) to remove impurities. The extracted solution from all fish was mixed and stored in individual aliquots of 0.75 ml in -20$^{o}$C.

Oxytocin treatment

To test the reversal of effects from non-functioning oxytocin signalling, *oxt* KO and WT controls were treated with either the fish homologue of oxytocin (isotocin; *Ser4*, *Ile8*-

Oxytocin; Cat. No. 4030890.0005, Bachem, Germany) or vehicle controls. For the treatment, first were anaesthetised by immersion in MS-222 solution (100mg/L), weighted and placed with their ventral part exposed in a pre-cut fissure on a spongy bed saturated in water. Fish were then administered a 2 µl/g ($\mu$ weight = 2.5g ± 0.8) intraperitoneal injection (30G needle) of either saline (vehicle control) or isotocin solution in saline (0.9%) at 1ng/kg, based on dose-response tests by Braida et al. [34]. The administration period was ~20s, after which fish were placed in a small compartment with tank water and allowed 2 min to recover with the help of oxygen supply from air bubbles slowly pipetted near their gills.

Behavioral test for social transmission of fear

Focal fish were randomly assigned to either of three conditions: to observe a shoal exposed to the alarm substance, to observe a shoal exposed to distilled water (vehicle control) or to be exposed to the alarm substance directly without observing any demonstrators (control for social transmission). The order of testing was randomized for each individual and conducted between 10:00 and 19:00. Animals were removed from home tanks on the day before experiments, randomly assigned to treatment groups, and kept in their experimental tank overnight for acclimatization. Experimental tanks had visual access to identical adjacent tanks (1.3 L; 12 × 12 × 15 cm), but were visually isolated from other external cues via opaque covers. Adjacent tanks kept either a shoal of two males and two females (Fig. 1A), or remained empty for the social transmission control. Each trial lasted for 15 min, including a 5 min baseline period followed by a 10 min post-exposure period. The alarm substance was kept on ice to avoid degradation during the trials and, thus, distilled water for the vehicle control was kept in the same conditions. Substances were administered via a flexible and transparent PVC tubing (diameter: 0.8 mm internal, 2.4 mm external).

Behavioral test of fear recognition

This test used video playbacks, which enabled us to control for inter-individual variation in demonstrators by presenting focal fish videos of the same fish in two separate states, neutral and fearful. The ability of fish to perceive and respond to videos of conspecifics under identical conditions, was demonstrated by Nunes et al. [33], which provide a detailed analysis of response to features of biological motion. Demonstrators used in video playbacks were recorded with a goPro camera (goPro hero3+, 60 fps, 1080 pixel resolution) placed in front of a 1.5 L tank, behind an opaque acrylic sheet with a customised cut-out for the camera lens, in order to keep the investigator covered during manipulations. The rest of the tank walls kept covered to reduce further visual interference and contained a flexible and transparent PVC tubing for substance administration (diameter: 0.8 mm internal, 2.4 mm external). A 10 minute recording of the tank was first captured to be used during the acclimatisation phase of experiments. Each fish used as a demonstrator was kept in the tank overnight, with the camera in place, and the following morning was recorded. During recording, following a 200s capture of baseline behaviour, the investigator released 0.75 ml (per 1.3L of water) of alarm substance and recorded for a further 200s, to capture erratic movement and freezing behaviour. Videos were edited using the VSDC© software (v. 6.3.6.18; Flash-Integro LLC, 2019) and included:

a 10 min video of the housing tank of demonstrators used as *background video* during acclimation (Fig. S); a 5 min *control video* with the demonstrator swimming (neutral state), which included 3 repetitions of a 100s swimming period (Video S1); a 5 min *stimulus video* with the demonstrator periodically exhibiting fear, which included 3 repetitions of a 60s swimming period followed by 40s bout of an erratic and freezing repertoire (Video S2). During tests, videos were displayed on monitors as real size images.

Experiments were carried out in a 4.5 L test tank (29.5 × 14.5 × 11 cm), stationed on a light box with infrared LEDs and with two LCD monitors (Asus VG248, 1080 HD, 144 Hz rapid refresh rate) positioned on either side, remaining visible through the glass walls **(**Fig. 4A**)**. The rest of the tank was covered with opaque lining and the overall set-up was housed in a compartment covered with a light-blocking black fabric to prevent visual interference from external stimuli. Playback screens were controlled and synchronised via a third screen connected to the same computer (TightVNC remote control software). Focal fish were kept in overnight isolation, housed individually in opaque tanks (12 × 12 × 15 cm) at 28 °C and a 14 L: 10 D photoperiod. The following day fish were individually placed in a central compartment of the test tank, devised by two removable transparent partitions, and acclimatized for 10m to the background video projected by both LCD screens. Following acclimation, one monitor was set to play the 5 min control video (neutral state) and the other the stimulus video (periodic fearful state), with the side of the video and the identity of the demonstrator (1 male and 1 female) counterbalanced across subjects. Following this stage, the partitions were lifted and fish allowed access to the entire tank for 10 min while playbacks on both sides displayed the control video (played twice in sequence).

Data extraction

For each test a continuous video-recording was obtained, using a high definition camera for the fear transmission test (Logitech B 525; acquisition at 30 fps) and a black-and-white camera with infrared sensitivity (Henelec 300B; acquisition at 30 fps) for the fear recognition test. The shift to the infra-red recording in the second test facilitated automated tracking over the larger arena, using the infra-red light backdrop from experimental set-up (Fig. 4A). Videos were fed to a remote laptop computer using the recording software Pinnacle Studio (v. 12, http://www.pinnaclesys.com). Individual recordings were then analyzed using the commercially available video tracking software Ethovision XT© 11.0 (Noldus Inc., The Netherlands).

For the fear transmission tests, recordings of the whole tank were tracked over the 15 min trial period in order to extract measures used to quantify fear behavior, both for the 5 min baseline period and the 10 min post-exposure period. These included proportion time spent exhibiting erratic movement [acceleration > 8 $cm/s^2$ and > 5 changes in direction/sec (> $90^{\circ}$)] and freezing (velocity < 0.2 cm/s).

For the fear recognition tests two separate stages were scored. First, for the first 5 min period of video observation, the zone of the central compartment in which animals were restricted was set as a region of interest (ROI) and animals tracked within this region. Attention to video playbacks was measured by the absolute compass heading (*x* direction relative to the stimulus video, ranging from 0° to 180°). Fear transmission was again measured by proportion time in erratic movement and freezing, and validated by added kinematic quantifiers [angular velocity (turn angle per frame); speed (cm/s)], and matched to the same measures extracted from the playback videos for validating transmission. Second, for the final 10 min of the test, during which full-tank access was allowed, the compartments next to each video were set as ROIs (Fig. 4A) and 3 measures were extracted: total distance travelled (exploration); latency time to first entry at either ROI (approach motivation); and the total time spend within each ROI (local preference).

Quantification of neuronal activation using the neuronal activation marker phospho-S6 ribosomal protein (pS6)

Brain tissue was collected for the quantification of brain activation and functional connectivity in *oxtr* experimental fish from the social transmission of fear experiment (1 hr post testing). Animals were anaesthetized with ice-cold water and their head extracted by cervical transection, fixed in 10% formalin for 3 days (in room temperature; RT), rinsed twice in 1× PBS (30min) and kept in EDTA (0.5 M, pH=8) for a further 2 days (RT). Coronal sections (5µm) of *oxtr* fish samples were extracted for immunohistochemical staining and microscopy, following paraffin-embedding.

Sectioned brains were stained for the pS6 . Slides were first kept in Tris-EDTA at 95°C (20 min) for antigen retrieval. Non-specific binding was blocked by a 1 hr in 1% BSA TBS incubation (0.025% Triton X-100) at RT and an overnight incubation in the primary antibody prepared in blocking solution (pS6 $Ser^{235/236}$ antibody D57.2.2E Rabbit mAB #4858 1:400; at 4 °C. Slides were then rinsed in TBS (0.025% Triton X-100) and incubated in the secondary antibody prepared in blocking solution (Alexa 594- Invitrogen goat anti-rabbit # A-11037 1:1000, Alexa 488- Invitrogen goat anti-chicken A-11039 1:1000). Slides were then washed in TBS with and then without 0.025% Triton X-100, before 20 min incubation in DAPI (4',6-diamidino-2-phenylindole) for nuclei counterstaining and rinsed in TBS before mounting (Biotium, Everbrite- 23003).

pS6 positive cells quantification projections visualization was performed on 20-fold magnified sections (Zeiss Axioscan.Z1 slide scanner) and analyzed via the Zeiss Zen blue 2.1 imaging software. Five consecutive coronal sections were quantified for each brain region (Fig. S5), where positive pS6 cells were counted in 1000 µm quadrants.

Imaging of oxytocin projections

OXT:GFP (Tg(*oxt*:EGFP)wz01 ID: ZDB-ALT-111103–1) positive fish were sacrificed in Tricaine, and their heads and skull removed. the heads were then fixed O.N at 4C in 4% PFA on a shaker. After fixation the brains were removed from the skull and subjected to whole-mount immunohistochemistry as per standard protocol: PFA was washed out, and samples were placed in ice cold (-20C) acetone in a freezer at -20C for 10 minutes. The acetone was washed out, and the samples were then incubated in blocking solution (PBS + 0.1% triton, 1% DMSO, 1% BSA, 5% NGS) for minimum of 2 hours at R.T and then incubated with primary Ab diluted at 1:200 ((anti-TH, Mouse monoclonal anti-Tyrosine hydroxylase, Merck-millipore, CAT: #MAB318) and anti-GFP (Chicken polyclonal anti-GFP IgY Antibody Fraction, Life Technologies, CAT: #A10262)) O.N at 4C on the shaker. The following morning, Samples were washed repeatedly (minimum of 6X15 minute washes) with blocking solution and then placed in blocking solution containing fluorescent secondary antibody at 1:200 O.N at 4C on a shaker. The following morning, Brains were submerged in 4% Noble Agarose, allowed to cool at 4C, and then sliced in a vibratome in ice-cold PBS at a thickness of 200 um. Slices were then mounted on a slide in mounting medium (Aqua-Polymount, polysciences, inc. 400 valley road, Warrington PA 18976, CAT: 18606-20) and imaged on a Zeis LSM 800 scanning confocal microscope.

Expression of oxytocin receptors in the brain

Brain tissue was also collected for testing the expression of oxytocin receptors (oxtr and oxtrl) in WT fish. WT fish samples were embedded in cryomoulds (OCT Compound, Tissue-Tek, Sakura 4583) and cryosectioned (150 µm coronal, Leica CM 3050S cryostat) for microdissection.

Receptor expression in WT fish was tested following microdissection of target brain areas from the cryosections, collected under stereoscope (Zeiss Stemi 2000) with a modified 210 µm needle (1 per region to prevent cross-contamination). Target areas were selected based on their involvement in social regulation and decision making [15] and included: the olfactory bulb (Ob), the medial zone of the dorsal telencephalic area (Dm, putative homologue of the mammalian basolateral amygdala), the preoptic area (POA), and the ventral nucleus of the ventral telencephalic area (Vv, putative homologue of the mammalian lateral septum), the dorsal nucleus of the ventral telencephalic area (Vd, putative homologue of the mammalian striatum), the supracommissural nucleus of the ventral telencephalic area (Vs, putative homologue of the mammalian medial extended amygdala and the bed nucleus of the stria terminalis), and the postcommissural nucleus of ventral telencephalic area (Vp). The Ob, Vv, Vd, Vs/Vp (pooled due to proximity), and POA were collected from both hemispheres at a single sampling point, due to their small size when compared to the diameter of the microdissection. The Dm was sampled from both hemispheres separately, and tissue was then pooled directly into lysis buffer and stored at -80°C until mRNA extraction.

For RNA extraction tissue was homogenized in 100 $\mu l$ of qiazol (lysis buffer) and incubated for 7 min at RT. 50 $\mu l$ of Chloroform was then added and shaken vigorously for 15 s and the sample left to incubate at RT for 5 min. Samples were then centrifuged at 13000 g for 20 min at 4ºC, and the upper aqueous phase transferred to a new tube where 1 volume of 70% ethanol was added. This mixture was transferred to an RNEasy® column and left to stand for 5 min at RT, and then was centrifuged for 1 min at 9000 g. A series of buffers from the

RNeasy® Lipid Tissue Mini Kit (Qiagen, 74804) were added to samples sequentially (700 *µl* of Buffer RW1, 500 *µl* of Buffer RPE and an additional 500 *µl* Buffer RPE), after each of which samples were centrifuged for 1 min at 9000 g and the flow-through discarded. The RNeasy column was then centrifuged a new 2 ml tube for 3 min at 14000 g and transferred to a separate new 1.5 ml tube where RNA was eluted with 25 *µl* of RNAse-free water, and centrifuged for 2 min at 9000 g. The elution step was repeated with the same RNAse-free water to increase RNA recovery efficiency and RNA screened for concentration, purity (260 nm and 280nm spectrophotometric absorbance; Thermo Scientific NanoDrop 2000) and integrity (Agilent 2100 Bioanalyzer).

Pooled RNA of each brain area was reverse transcribed to cDNA (iScript cDNA Synthesis Kit, Biorad, 1708890) following the manufacturer's instructions. Briefly, in a clean Eppendorf tube, nuclease-free water, 5x iScript reaction mix (4*µl*), iScript reverse transcriptase (1*µl*), and RNA template (100 fg to 1 *µg* total RNA) were added up to a total volume of 20 *µl* and incubated in a PCR thermocycler (5 min priming at 25°C, 60 min reverse transcription at 42°C, 5 mins reverse transcription inactivation at 85°C, and kept at 4°C). Samples were subsequently stored in -20°C until use.

Diluted cDNA samples (1:10) were used as templates for quantitative polymerase chain reactions (qRT-PCR). Primer sequences for the oxytocin receptor (*oxtr*) and the reference gene (*eef1a1l1*: eukaryotic translation elongation factor 1 alpha 1, like 1) were designed in the Primer 3 software (Premier Biosoft International, Palo Alto, CA, USA) and primer sequences for oxytocin receptor-like (*oxtrl*) were provided by Gil Levkowitz. qRT-PCR reactions were performed in the Applied Biosystems quantstudio 7 thermocycler (7900 HT, Thermofisher) in 8 *µl* triplicate reactions with SYBR Green PCR Master Mix (Applied Biosystems, Thermofisher) with 50 µM primers for *oxtr* and *eef1a*, and 13.3 µM for *oxtrl*. Thermocycling conditions were 5 min at 95°C, followed by 40 cycles of 95°C for 30 s, annealing temperature 60°C for 30 s, and extension at 72°C for 30 s. After PCR, a melting curve program from 55 to 95°C with 0.5°C changes was applied and fluorescence cycle thresholds (*Ct*) were automatically measured.

### Analysis

Statistical analyses, calculations and graphical representations were carried out using the software Graphpad Prism® (v. 8.0.1; GraphPad LLC, San Diego, CA), R© (v. 4.0.3; R Core Team) and Minitab® (v.17; Minitab Inc., State College, PA). Figures were edited and completed with illustrations using the software Adobe® Illustrator® (CS6, v.16.0.0; Adobe Systems Inc.) and Inkscape© (v. 0.92.4; Free Software Foundation Inc.). Continuous data were tested for normality using the Ryan-Joiner and Kolmogorov-Smirnov tests and homogeneity of variance was tested using the Bonnett's and Levene's tests. Finite ranging and proportion based scores were tested for normality using the D'Agostino & Pearson test ($K^2$), which confirms Gaussian distribution via skewness and kurtosis [35].

*Genetic expression of receptor genes* was calculated using the $2^{-\Delta Ct}$ method [36]:

$$2^{-\Delta Ct} = 2^{Ct_{Ref} - Ct_{Target}}$$

where $Ct_{Ref}$ is the cycle threshold for the reference gene and $Ct_{Target}$ is the cycle threshold for the target gene. Therefore, target gene expression was represented as relatively to the reference gene and quantified by the mean of this value across three technical replicates.

*Cell counts of pS6 positive cells* were tested via a generalized linear model with quasi-Poisson regression, with treatment (alarm substance or control) and genotype as fixed factors. A backward stepwise procedure was used to exclude non-significant effects, followed by *post-hoc* comparisons corrected with the two-stage linear step-up procedure for FDR-adjusted p-values.

*Percentage time erratic and freezing* were compared between treatment groups using unpaired *t*-tests for parametric data, Welch's 2-sample *t*-tests for non-homogeneous normal data, and Mann-Whitney *U* tests for non-parametric data. Comparisons between lines were carried out using ANOVA tests, either at the two-way with treatment (alarm or control), or at the three-way when examining added effects from oxytocin injection (versus control) for testing recovery. Data not conforming to parametric assumptions were log-transformed $[log_{10}(x + 1)]$. *Post-hoc* comparisons for testing differences between the treatments were corrected with False Discovery Rate (FDR) and *p*-value adjusted (Benjamini and Hochberg's method).

*Compass orientation* (absolute heading in degrees) was tested for differences from 90˚ by 1-sample *t*-tests to compare deviations from divided attention (see Fig. 2A) and compared between groups using Welch's 2-sample *t*-tests (due to unequal sample sizes).

*Angular velocity and speed* for each individual was measure by mean values from across the 3 replicates during observation, for both the 40s stress demonstration period and separately for the immediately preceding 40s under control conditions (neutral, swimming), in 10s time bins. For each of the two periods, stress and neutral control demonstration, we calculated the total area under the curve (AUC) across time bins using the trapezoid approximation method:

$$AUC_{t_1 - t_2} = \frac{(x_1 + x_2)}{2} \times (t_1 - t_2)$$

Behavioural change between states was then calculated as the difference in AUC between the control and stress demonstration periods ($\Delta$ AUC), for both speed and angular velocity. These values of change were tested for significant deviation from no difference ($\mu \neq 0$) by 1-sample *t*-tests and compared between groups using Welch's 2-sample *t*-tests (due to unequal sample sizes). To examine the degree of stress contagion, we tested consistency between observer and demonstrator angular velocity and speed using a linear regression model with time bin as an interaction term for time-dependent changes.

*Total distance travelled* (cm) was compared between genotypes for all lines by Welch's 2-sample *t*-tests. To test for the effect of injections we used a two-way ANOVA with treatment (oxytocin or vehicle injection) and its interaction with genotype (WT or *oxt* KO) as predictors, and *post hoc* comparisons using Fisher's LSD.

*Approach latency* (s) towards the stimulus video ROI, where the demonstrator periodically stressed during observation, was compared to the approach latency towards the control-video ROI by Welch's 2-sample *t*-tests on the mean (due to unequal sample sizes) and effect sizes calculated using Cohen's *d* and the proportion of mean change, for each group. To assess the reversal of effects following injection treatments, for both the oxytocin and the vehicle treatment, we compared the directional effect size between WT and *oxt* KO. To do this we first calculated the sampling variance of each group:

$$v = \frac{1}{n_1} + \frac{1}{n_2} + \frac{d^2}{2(n_1 + n_2)}$$

and used this to compare Cohen's *d* values between genotypes, using a *z*-test:

$$z = \frac{d_1 - d_2}{\sqrt{v_1 - v_2}}$$

where for normal distributions the null hypothesis ($H_0 : d_1 = d_2$) can be rejected if $|z| \geq 1.96$ at $\alpha = 0.05$ (two-sided).

*Preference scores* (PS) were calculated based on the time spend in ROI near the stimulus ($ROI_S$) and the control video ($ROI_C$) using:

$$PS = \frac{(ROI_S - ROI_C)}{(ROI_S + ROI_C)}$$

where values range between -1 (full preference for control) and 1 (full preference for stimulus). In order to validate that preference levels were statistically different from chance, we first tested if mean preference scores for each group were significantly greater or lower than 0, depending on the direction, by using 1-sample *t*-tests. Comparisons of preference scores between WT and KO fish for each line were performed using Welch's 2-sample *t*-tests (due to unequal sample sizes). To test for the effect of injections we used a two-way ANOVA with treatment (oxytocin or vehicle injection) and its interaction with genotype as predictors, and *post hoc* comparisons using Fisher's LSD.

Brain network analysis

We constructed networks representing the co-activation patterns for each treatment as follows. For the case of M specimens, characterized by one sample (cell count) reading $x_i$ for each of the N brain regions, one typically considers the M-dimensional vector $\mathbf{x}_i$, where i labels the region, and then computes the correlations $c_{ij}$ for all region pairs, obtaining a weighted correlation matrix that we interpret as the adjacency matrix of a functional network. We then filter each layer using the networks for each treatment as described by De Vico Fallani et al. [37] keeping the links with larger weight (in absolute value) up to a threshold density of $\rho = 0.17$. For the analysis of the excitation and inhibition pattenrs, we separate the weighted and signed graph in two subgraphs containing respectively only the positive and negative links, which we interpret as pertaining to network excitation and inhibition configurations. Distributions were compared by the two-sample Kolmogorov-Smirnov test and averages using the Mann-Whitney *U* test.

*For the ranking of node strengths and correlations* we compared, for all treatments, the strength (weighted degree of the full correlation matrix) of nodes in the standard way and rank them in descending order. To assess, similarities between ranking corresponding to different treatments, we use the Kendall-Tau rank correlation and retain as significantly difference from zero only correlations with p<0.01.

*For the Detection of communities and extraction of conserved submodules* we used the spin glass community detection method [38] applied on the average treatment network, obtained by averaging over the corresponding graph tower matrices. To increase the robustness of the detection, for each treatment, we repeated the community detection 1000 times. We computed the (center) consensus partition for each treatment from the 1000 candidate partitions we extracted as described by Peixoto [39].

To identify parts of modules that are shared between communities in different treatments, we study the distribution of intersection sizes. In particular, for two partitions consider partitions $P_y = \{C_y^0, C_x^1, \dots.. C_x^m\}$ and $P_y = \{C_y^0, C_y^1, \dots.. C_y^l\}$ for treatments x and y, and then compare for each pair of modules $(C_x^i, C_y^j)$ the intersection $J_{x,y}^{i,j} = C_x^i \cap C_y^j$ and then measure its cardinality $|J_{x,y}^{i,j}|$. We compute the significance of the measured intersection sizes by a permutation test based on a null distribution $p_0(|J|)$ constructed as follows: for a pair $(C_x^i, C_y^j)$, we sample uniformly at random 10000 pairs of node sets with cardinality respectively $|C_x^i|$ and $|C_y^j|$and compute the size $|J|$ of their intersection. We consider statistically significant and thus retain the observed submodules $J_{x,y}^{i,j}$ such that $|J_{x,y}^{i,j}| > \mu(|J|) + 3\sigma(|J|)$ (equivalent to a p < 0.01 significance threshold), where $\mu(|J|)$ and $\sigma(|J|)$ are the first two moments of $p_0(|J|)$.

| Table S1. Ranks of node strengths for each treatment for both wild-types and mutants. | | | | |
|---|---|---|---|---|
| | **Wild-types** | | **Mutants** | |
| | **Control** | **Alarm** | **Control** | **Alarm** |
| **0** | DL | PPP | PM | VP |
| **1** | DP | VS | OB | VV |
| **2** | PPAM | PPAL | PPP | PPAL |
| **3** | DM | DL | DM | VS |
| **4** | PPAL | PPAM | HAV | PPP |
| **5** | ATN | DP | VV | HAD |
| **6** | PPP | OB | ATN | D |
| **7** | PM | VP | PPAM | OB |
| **8** | OB | VV | DL | ATN |
| **9** | VP | PM | DP | LH |
| **10** | D | D | VP | DM |
| **11** | VD | HAV | PPAL | DP |
| **12** | LH | ATN | VC | DL |
| **13** | VC | DM | D | HV |
| **14** | HV | VD | HV | PPAM |
| **15** | HAV | VC | HAD | HAV |
| **16** | VS | LH | VD | VD |
| **17** | VV | HAD | LH | PM |

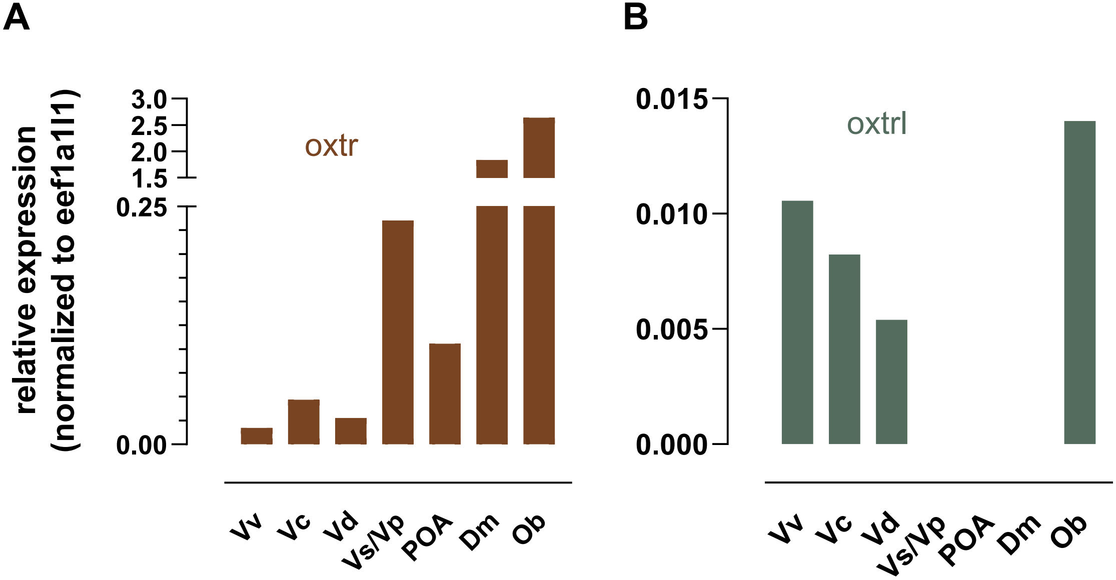
A
B
relative expression
(normalized to eef1a1l1)
3.0
2.5
2.0
1.5
0.25
0.00
oxtr
Vv
Vc
Vd
Vs/Vp
POA
Dm
Ob
0.015
0.010
0.005
0.000
oxtrl
Vv
Vc
Vd
Vs/Vp
POA
Dm
Ob

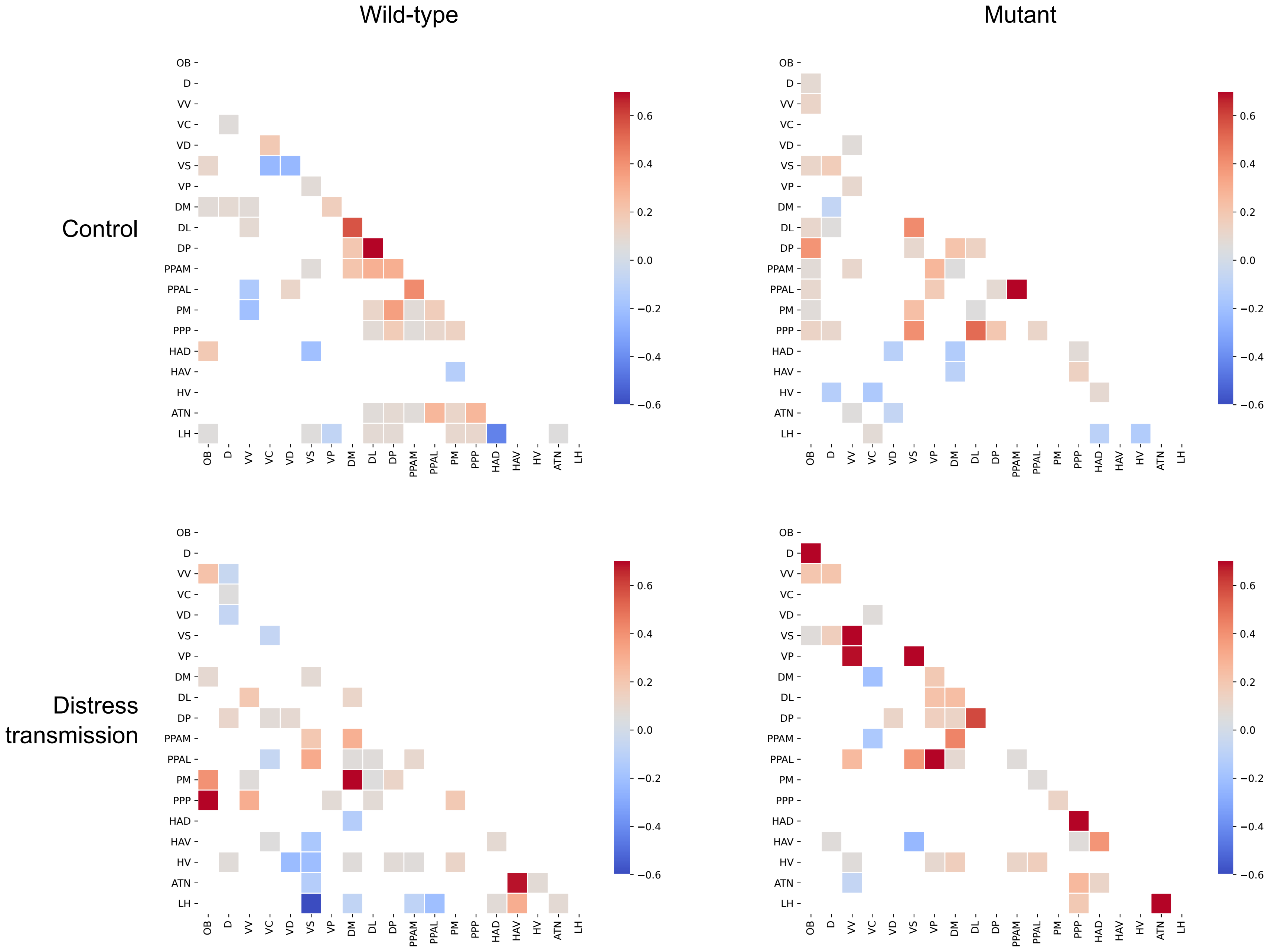

Wild-type
Mutant
Control
Distress transmission
OB
D
VV
VC
VD
VS
VP
DM
DL
DP
PPAM
PPAL
PM
PPP
HAD
HAV
HV
ATN
LH
0.6
0.4
0.2
0.0
−0.2
−0.4
−0.6

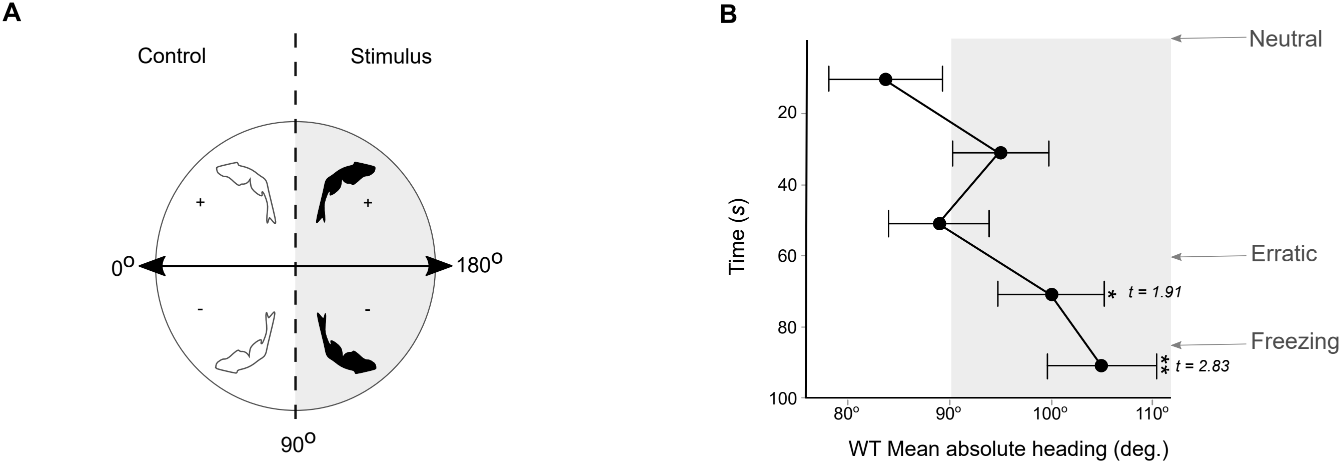

A
Control
Stimulus
+
+
-
-
0o
180o
90o
B
Neutral
Erratic
Freezing
20
40
60
80
100
Time (s)
80o
90o
100o
110o
WT Mean absolute heading (deg.)
*
t = 1.91
**
t = 2.83

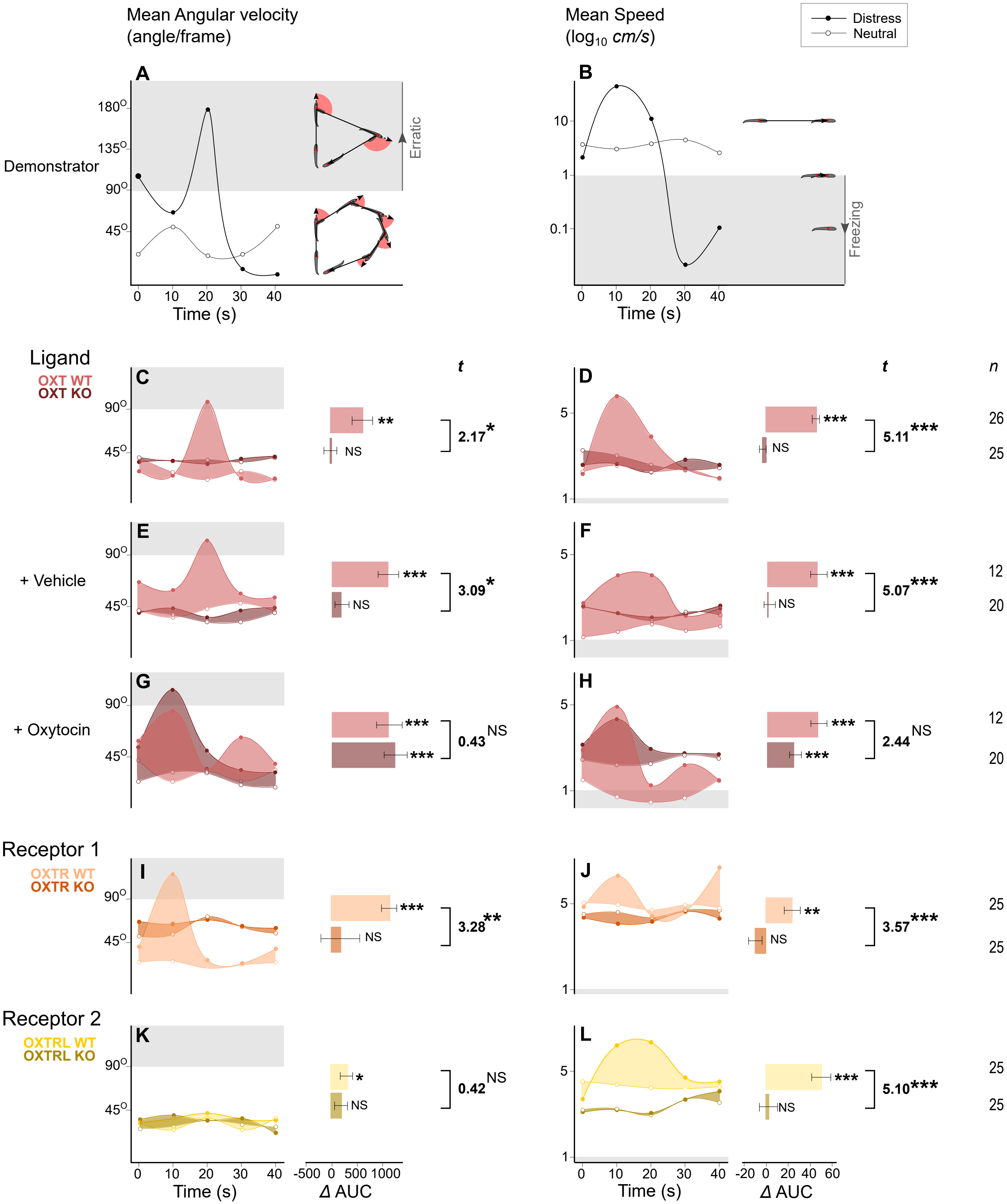

Mean Angular velocity (angle/frame)
Mean Speed ($log_{10}$ *cm/s*)
Distress
Neutral
A
B
Demonstrator
Erratic
Freezing
Time (s)
Ligand
OXT WT
OXT KO
+ Vehicle
+ Oxytocin
Receptor 1
OXTR WT
OXTR KO
Receptor 2
OXTRL WT
OXTRL KO
t
n
2.17*
3.09*
0.43 NS
3.28**
0.42 NS
5.11***
5.07***
2.44 NS
3.57***
5.10***
26
25
12
20
12
20
25
25
25
25
Δ AUC
